# Red blood cell tension controls *Plasmodium falciparum* invasion and protects against severe malaria in the Dantu blood group

**DOI:** 10.1101/475442

**Authors:** Silvia N. Kariuki, Alejandro Marin-Menendez, Viola Introini, Benjamin J. Ravenhill, Yen-Chun Lin, Alex Macharia, Johnstone Makale, Metrine Tendwa, Wilfred Nyamu, Jurij Kotar, Manuela Carrasquilla, J. Alexandra Rowe, Kirk Rockett, Dominic Kwiatkowski, Michael P. Weekes, Pietro Cicuta, Thomas N. Williams, Julian C. Rayner

**Affiliations:** KEMRI-Wellcome Trust Research Programme, Kilifi, Kenya; Wellcome Sanger Institute, Cambridge, UK; Cavendish Laboratory, University of Cambridge, Cambridge, UK; Cambridge Institute for Medical Research, Cambridge, UK; University of Edinburgh, Edinburgh, UK; Wellcome Centre for Human Genetics, University of Oxford, Oxford, UK; Big Data Institute, University of Oxford, Oxford, UK; Imperial College London, London, UK

## Abstract

Malaria has had a major effect on the human genome, with many protective polymorphisms such as sickle cell trait having been selected to high frequencies in malaria endemic regions^1, 2^. Recently, we showed that a novel blood group variant, Dantu, provides 74% protection against all forms of severe malaria in homozygous individuals^3-5^. This is a similar degree of protection to sickle cell trait and considerably greater than the most advanced malaria vaccine, but until now the mechanism of protection has been unknown. In the current study, we demonstrate a significant impact of Dantu on the ability of *Plasmodium falciparum* merozoites to invade RBCs. The Dantu variant was associated with extensive changes to the RBC surface protein repertoire, but unexpectedly the malaria protective effect did not correlate with specific RBC-parasite receptor-ligand interactions. By following invasion using video microscopy, we found a strong link between RBC tension and parasite invasion and, even in non-Dantu RBCs, identified a tension threshold above which RBC invasion did not occur. Dantu RBCs had higher average tension, meaning that a higher proportion of Dantu RBCs could not be invaded. These findings not only provide an explanation for the protective effect of Dantu against severe malaria, but also provide fresh insights into the essential process of *P. falciparum* parasite invasion, and how invasion efficiency varies across the heterogenous populations of RBCs that are present both within and between individuals.

---

The Dantu polymorphism has been fine mapped to a structural rearrangement in the glycophorin (*GYP*) gene cluster. This rearrangement of the *GYPA* and *GYPB* genes creates two copies of a hybrid gene that encodes the Dantu blood group antigen, a novel sialoglycoprotein composed of a glycophorin B (GYPB) extracellular domain fused with a glycophorin A (GYPA) intracellular domain^5^. Both GYPA and GYPB are known to play an important functional role in the invasion of *P. falciparum* merozoites into RBCs: GYPA is the receptor for the *P. falciparum* erythrocyte-binding ligand *Pf*EBA-175^6^, while GYPB is a receptor for *P. falciparum* erythrocyte binding ligand 1 (*Pf*EBL-1)^7^.

To investigate the impact of Dantu on *P. falciparum* invasion, we collected RBC samples from 42 healthy children from the Indian Ocean coastal region of Kenya, where this polymorphism is found at a minor allele frequency of approximately 10% - the highest frequency yet described^3^. To eliminate any possible confounding from other large-effect malaria protective polymorphisms, we only included samples from subjects who were negative for both sickle cell trait and homozygous α-thalassaemia (**Supplementary Table 1**). We employed a previously described flow-cytometry-based preference invasion assay to quantify invasion^8^. Parasites were co-cultured with differentially labelled Dantu-heterozygous, homozygous, and non-Dantu RBCs, and we measured invasion events using a fluorescent DNA dye. Three parasite strains, 3D7, Dd2 and SAO75, showed a significantly reduced ability to invade Dantu RBCs, with a similar trend observed for GB4 and 7G8. These five strains were chosen for their use of a variety of invasion pathways and their differing reliance on sialic-acid dependent receptors such as glycophorins^9^; Dantu appears to limit invasion in all cases. Dantu homozygotes were more resistant to invasion than heterozygotes, suggesting a dose-dependent effect (**Fig. 1a, Supplementary Table 2**).

**Figure 1.**
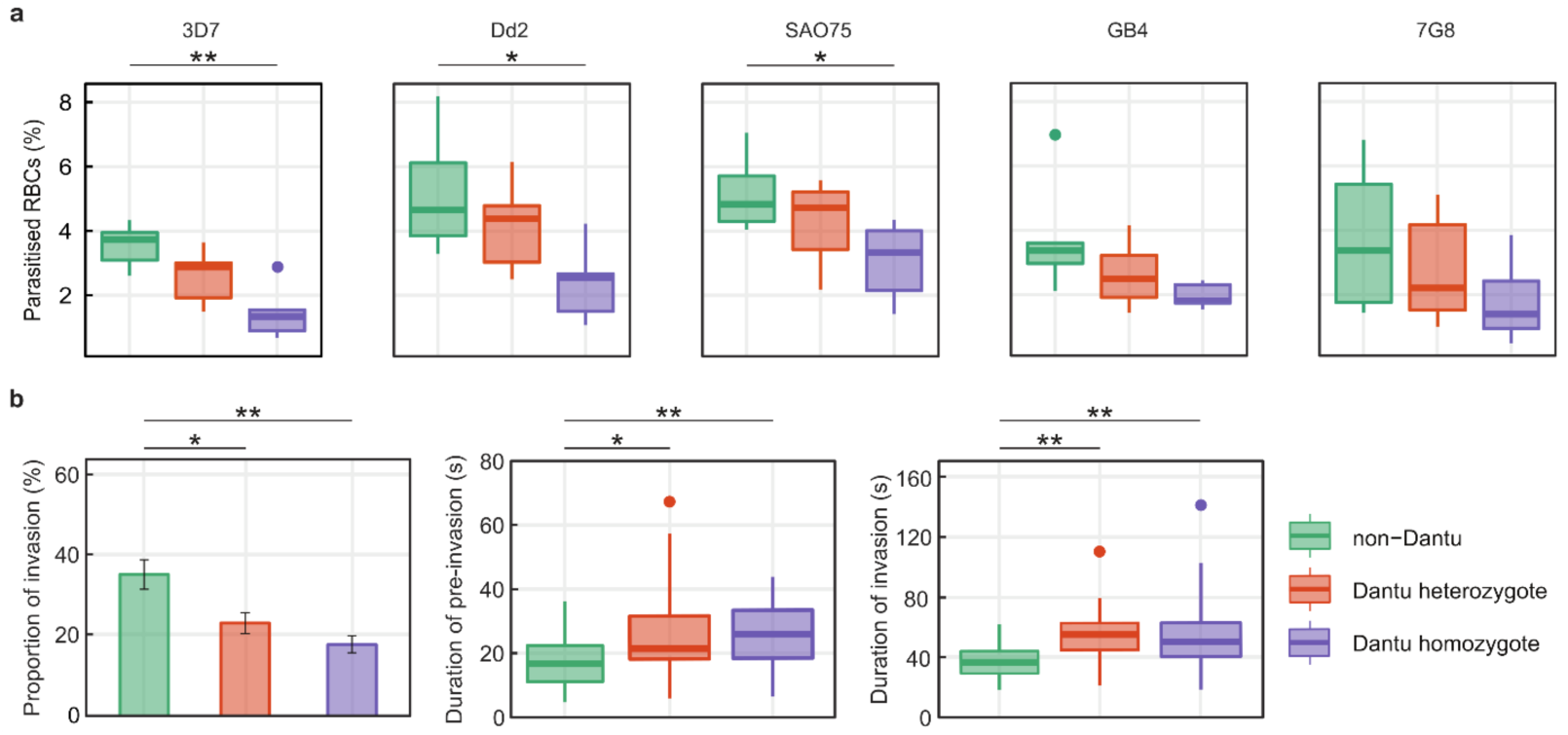
Reduced invasion of Dantu variant RBCs by multiple *Plasmodium falciparum* strains. **(a)** The relative ability of *P. falciparum* strains from multiple geographic locations (3D7 and GB4 West Africa; Dd2 Southeast Asia; SA075 East Africa; 7G8 Brazil) to invade RBCs from each Dantu genotype group was measured using a flow cytometry-based preference invasion assay. The percentage of parasitised RBCs that successfully invaded each genotype group is indicated on the y-axis. (**b**) The invasion process was also follow by live video microscopy, where the invasion rate of 3D7 merozoites into RBCs in each genotype group is measured as the proportion of merozoites that contacted and successfully invaded RBCs, relative to all merozoites that contacted RBCs. Pre-invasion time lasts from the first merozoite contact through RBC membrane deformation and resting, while invasion time is from the beginning to the end of merozoite internalization. Six RBC samples per genotype group were tested in both the flow and video microscopy assays. Boxplots indicate the median (middle line) and interquartile ranges (top and bottom of boxes) of the data, while whiskers denote the total data range. The barplot shows the mean and standard deviation of the live video microscopy invasion data. Statistical comparison across the three genotype groups was performed using a one-way ANOVA test, while pairwise comparisons between genotype groups was performed using the Tukey HSD test and Mann-Whitney U test. **p < 0.01; *p < 0.05.

To investigate the specific step at which invasion was impaired, we studied the invasion process for 3D7 strain parasites using time-lapse video microscopy. Invasion into Dantu RBCs was also significantly decreased in this real-time assay, validating the results of our FACS-based assays (**Fig. 1b and Supplementary Video Material**). Even when successful, the kinetics of invasion into Dantu homozygous RBCs was affected, with both the early pre-invasion step and the subsequent active invasion step being significantly longer (**Fig. 1b**). By contrast, we found no significant difference in the strength of attachment between merozoites and Dantu RBCs, as measured using optical tweezers^10^ (**Supplementary Fig. 2**), nor was there a significant difference in the degree of membrane deformation^11^ or echinocytosis triggered by parasites invading RBCs of differing Dantu genotype (**Supplementary Fig. 1**). Overall, these results indicate that Dantu has a pleiotropic effect on invasion across both contact and entry of merozoites into RBCs.

**Figure 2.**
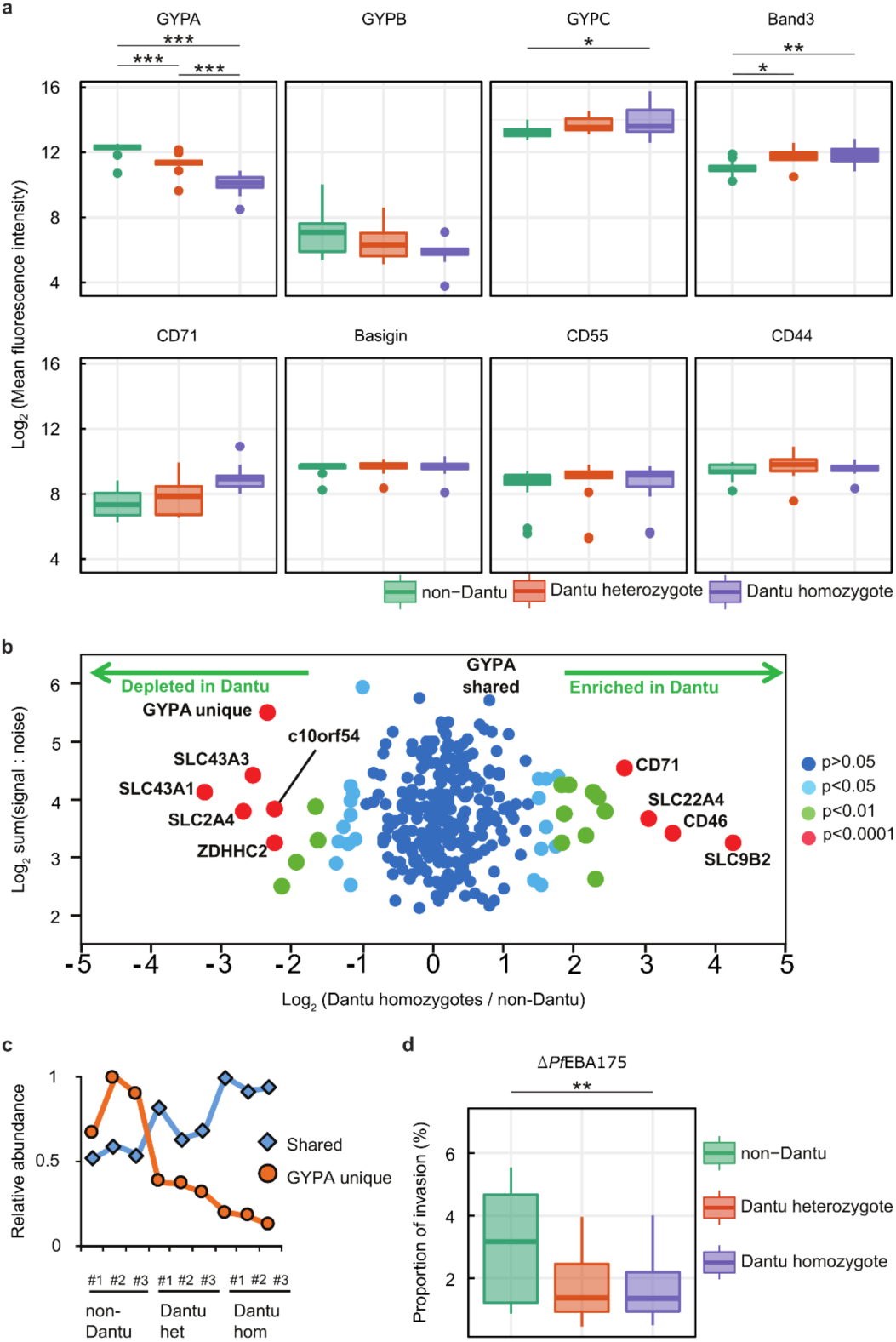
RBC membrane protein characteristics vary across Dantu genotype groups but do not directly correlate with invasion efficiency. (**a**) The relative expression of essential RBC membrane proteins across genotype groups was assessed using fluorescent monoclonal antibodies in flow cytometry assays. 13 non-Dantu, 12 Dantu heterozygotes and 11 Dantu homozygote RBC samples were tested. (**b**) Scatter plot of all proteins quantified by mass spectrometry. Fold change was calculated by average signal:noise (Dantu homozygotes) / average signal:noise (non-Dantu) (3 samples per genotype group). The GYPA protein was split into two parts: identified by peptides unique to GYPA (‘GYPA unique’, all originating from the extracellular region) or identified by peptides shared with the Dantu protein (‘GYPA shared’, all originating from the intracellular region). Benjamini-Hochberg-corrected significance was used to estimate p-values. (**c**) Graph of the relative abundance of ‘unique’ and ‘shared’ GYPA peptides across all donors. Signal:noise values were normalised to a maximum of 1 for each protein. (**d**) The invasion efficiency of a genetically modified parasite strain, Δ*Pf*EBA175, was compared across genotype groups (13 non-Dantu, 12 Dantu heterozygotes and 12 Dantu homozygote RBC samples) using the flow-cytometry-based preference invasion assay. Boxplots indicate the median (middle line) and interquartile ranges (top and bottom of boxes) of the data, while whiskers denote the total data range. Statistical comparison across the three genotype groups was performed using a one-way ANOVA test, while all pairwise comparisons between genotype groups was performed using the Tukey HSD test. ** p < 0.01; * p < 0.05

Comparing haematological indices across the Dantu genotypic groups revealed significantly lower mean cell volumes (MCVs) and mean cell haemoglobin concentrations in RBCs from Dantu homozygotes (**Table 1**). This suggests that Dantu directly impacts RBC properties, perhaps by altering the composition of the RBC surface. Despite the fact that the Dantu polymorphism leaves an intact copy of the *GYPA* gene within the genome^5^, the surface expression level of GYPA was significantly reduced in Dantu RBCs when measured by flow cytometry (**Fig. 2a**), an observation which confirms that made in previous small-scale studies^12^. By contrast, surface expression of GYPB was unchanged, while both GYPC, another important invasion receptor^13^, and Band 3 were increased in Dantu RBCs (**Fig. 2a**). We also observed a higher surface expression level of CD71, a marker of younger stages of RBC development that is lost as RBCs mature (**Fig. 2a**). We saw no differences in the surface expression levels of other RBC membrane proteins involved in parasite invasion, including Basigin, CD55 and CD44 (**Fig. 2a**).

**Table 1.** Clinical and demographic characteristics of study participants

| Characteristic | non-Dantu | Dantu Heterozygotes | Dantu Homozygotes | P value | P adj |
| --- | --- | --- | --- | --- | --- |
| Mean Age (SD; years) | 11.4 (1.6) | 11.6 (1.7) | 9.3 (4.1) | 0.255 | - |
| Sex (F/M) | 5/12 | 5/11 | 5/8 | 0.866 | - |
| Red blood cell count (SD; $10^6/\mu\text{L}$) | 4.60 (0.36) | 4.62 (0.34) | 4.94 (0.68) | 0.252 | 0.286 |
| Reticulocyte count (SD; $10^6/\mu\text{L}$) | 0.02 (0.01) | 0.04 (0.01) | 0.05 (0.02) | 0.009 | 0.545 |
| White blood cell count (SD; $10^3/\mu\text{L}$) | 5.83 (1.46) | 5.67 (0.78) | 6.89 (2.93) | 0.433 | 0.643 |
| Platelet count (SD; $10^3/\mu\text{L}$) | 284.06 (77.91) | 293.31 (73.91) | 346.92 (96.40) | 0.122 | 0.106 |
| Haematocrit (SD; %) | 37.23 (1.66) | 37.23 (2.68) | 36.22 (2.42) | 0.279 | 0.188 |
| Haemoglobin concentration (SD; g/dL) | 12.54 (0.54) | 12.59 (0.93) | 12.08 (1.13) | 0.270 | 0.168 |
| Mean cell volume (SD; fL) | 81.32 (4.69) | 80.78 (4.67) | 74.40 (9.66) | 0.082 | 0.015* |
| Mean cell haemoglobin (SD; pg) | 27.38 (1.75) | 27.31 (1.83) | 24.90 (4.12) | 0.119 | 0.015* |
| Mean cell haemoglobin concentration (SD; g/dL) | 33.67 (0.60) | 33.81 (0.63) | 33.33 (1.51) | 0.888 | 0.113 |
| Red blood cell distribution width (SD; %) | 12.83 (1.29) | 12.70 (1.13) | 13.78 (1.66) | 0.161 | 0.830 |

Data are mean values, with the standard deviation (SD) in parentheses. Statistical comparison across the three genotype groups was performed using the Kruskal-Wallis test. P adj = p-values adjusted for age and sex of the participants. \*p < 0.05.

To quantify these changes more accurately, we employed plasma membrane profiling^14, 15^, which uses membrane-impermeable oxidation and aminooxybiotinylation in combination with tandem mass tag mass spectrometry to quantify surface proteins. Analyzing RBCs from three donors of each type allowed us to quantify 294 proteins that were either anchored in the RBC membrane or had a transmembrane region. The data revealed widespread cell surface changes, with 40 proteins up-regulated and 35 down-regulated by >50% in Dantu heterozygotes, and 66 proteins up-regulated and 35 down-regulated by >50% in Dantu homozygotes (**Figures 2b, Supplementary Fig. 3, Supplementary Table 6**). Significant changes in GYPA and Band 3 were identified, in keeping with our flow cytometry data. Mass spectrometry was also able to distinguish between peptides unique to GYPA (which were all in the extracellular region of the protein) from those shared with Dantu (which were all intracellular), confirming the presence of the Dantu antigen on the RBC surface (**Figure 2c**).

**Figure 3.**
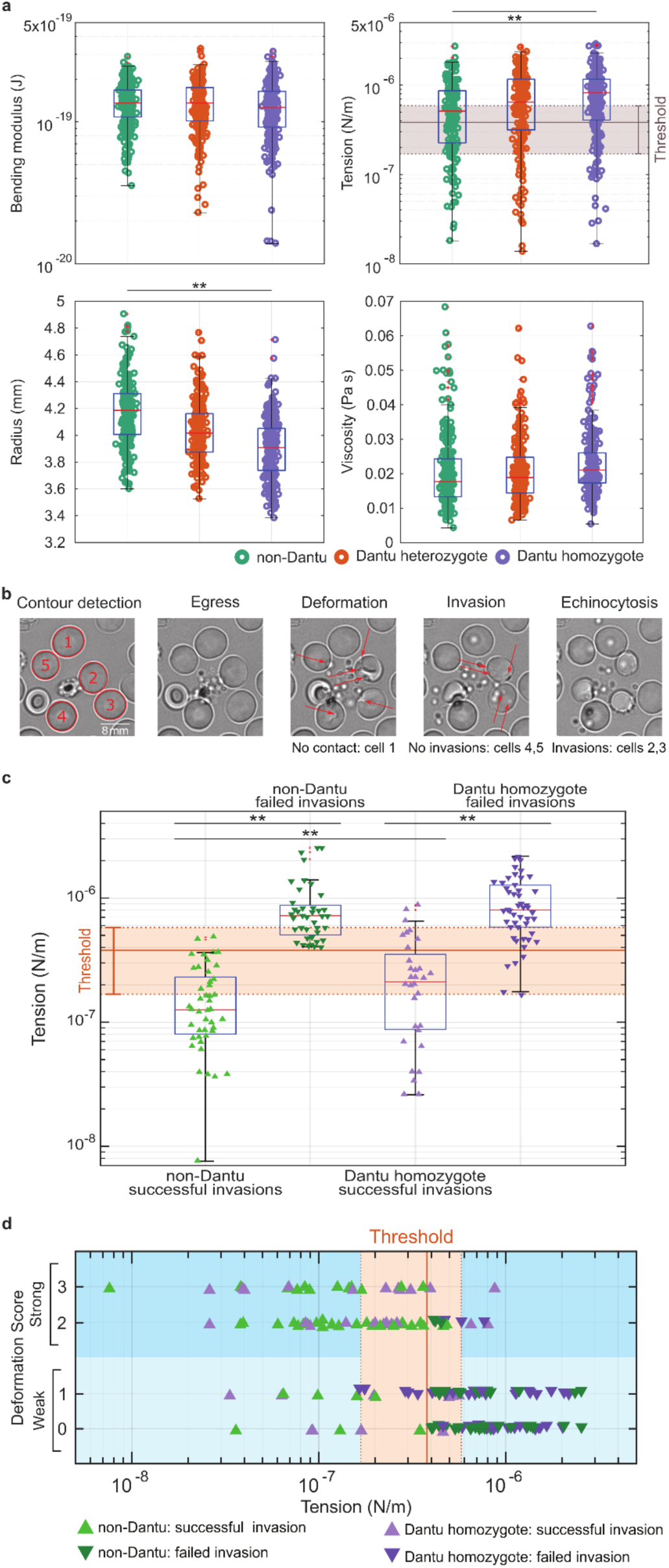
Biomechanical properties of the RBC membrane differ across Dantu genotypes and correlate with invasion. (**a**) Membrane flickering spectrometry enabled measurement and comparison of RBC bending modulus, tension, radius, and viscosity across Dantu genotype groups (6 samples per genotype group). Mean and standard deviation for tension were (6.0 ±2.8) *10^−7^ N/m in non-Dantu, (7.9±2.8) *10-7 N/m in Dantu heterozygotes, and (8.8 ± 0.7) *10^−7^ N/m in Dantu homozygotes. The impact of tension on parasite invasion was evaluated by simultaneously measuring the tension from flickering analysis and imaging the invasion process from rupturing schizonts (“egress”, “deformation”, then either “invasion” and “echinocytosis”, or a failed invasion) in live video microscopy (**b**), and experiments were done on RBCs from non-Dantu and Dantu homozygous individuals (**c**). The impact of tension on RBC deformation during pre-invasion, induced by merozoites contacting RBCs, was compared across Dantu genotype groups (**d**). The threshold range for tension, marked in (a), (c) and (d) was obtained by comparing distributions of tension across Dantu genotypes with their invasion efficiency. Deformation scores were classified as first described in ^11^. Pairwise comparisons between genotype groups was performed using the Mann-Whitney U test. ** p < 0.01. Number of cells and tension reported in Supplementary Tables 3, 4.

Two independent methods therefore confirm that there are major changes in the surface of Dantu RBCs, including the known *P. falciparum* invasion receptor, GYPA, which was significantly reduced. To test whether these changes might explain the impact of Dantu on RBC invasion, we investigated the invasion efficiency of a genetically modified *P. falciparum* strain that had *Pf*EBA175, the ligand for GYPA, disrupted^16^. Surprisingly, we observed a significant reduction in the ability of Δ*Pf*EBA175 parasites to invade Dantu RBCs (**Fig. 2d, Supplementary Table 2**). Given that this parasite cannot, by definition, use GYPA for invasion, changes in the receptor-ligand interaction between GYPA and *Pf*EBA175 cannot alone explain the inhibitory impact of Dantu.

The Dantu polymorphism therefore affects invasion across multiple and diverse *P. falciparum* strains, regardless of their reliance on GYPA as an invasion receptor, and impacts multiple steps in the invasion process, including the earliest steps before GYPA is thought to be involved^17^. All these observations support the conclusion that the protective effect of Dantu is not related directly to receptor-ligand interactions but might instead relate to its broader RBC biomechanical effects. Measuring membrane contour fluctuations using live imaging and flickering spectrometry^18^ allowed us to generate direct measurements of RBC membrane mechanics, such as tension, radius, viscosity, and bending modulus, across genotype groups. These phenotypes vary naturally between RBCs, which can circulate for more than three months after they are produced from stem cells. Dantu RBCs had significantly higher tension and a smaller radius than non-Dantu RBCs, as expected due to their lower MCV, whereas no significant differences were seen in bending modulus and viscosity (**Fig. 3a**). Tension and radius are linked properties, with higher tension leading directly to smaller RBCs^19^.

To test whether there was any link between tension and invasion, we measured both parameters simultaneously using video microscopy in separate experiments for non-Dantu or Dantu variant RBCs. Tension was measured for all cells adjacent to a rupturing schizont using high frame rate capture, then the fate of invasion of all parasite-RBC contact events into the same cells was monitored following schizont rupture (**Fig. 3b**). We observed an intrinsic distribution of membrane tensions in RBC from each donor, which led us to discover a clear association between tension and invasion: merozoites preferentially invaded neighbouring RBCs with low tension (**Fig. 3c, Supplementary Table 4**). Comparing the distributions of tension for Dantu and non-Dantu RBCs with the percentage of successful invasions into these cells showed that there was a tension threshold for successful parasite invasion – 3.8 (±2.0) *10^−7^ N/m. Successful invasion, both in Dantu and non-Dantu RBCs, was very rare above this threshold (**Fig. 3c)**. There was also an association between tension and deformation during the invasion process, with RBCs above the tension threshold only ever weakly deformed (scores 0 and 1), whereas RBCs with tensions below the threshold were more strongly deformed (scores 2 and 3), as well as being more frequently invaded by parasites (**Fig. 3d, Supplementary Table 4**). While osmotic stress, which influences tension, has been broadly related to invasion efficiency before^20^, and chemically-induced increase in tension has been shown to reduce invasion^21^, this is the first time that tension and invasion have been followed at single-cell single-event resolution, thereby proposing and measuring a specific tension threshold for invasion. Critically, the median tension of Dantu RBCs was 8.4*10^−7^ N/m, which means that the majority of Dantu RBCs are above the tension threshold of 3.8 (±2.0) *10^−7^ N/m (**Fig. 3a**). The impact of Dantu on the biomechanical properties of the RBC is therefore sufficient to explain its impact on invasion, and is in keeping with the invasion-inhibitory effect of Dantu being consistent across multiple *P. falciparum* strains, regardless of their reliance on GYPs or other invasion receptors.

In summary, we have established a mechanism of protection for a recently identified complex structural polymorphism with strong protective effects against severe malaria. We demonstrate a marked inhibitory impact of the Dantu polymorphism on *P. falciparum* RBC invasion, and establish that while there is an effect of Dantu on the RBC surface expression level of multiple proteins, it is its impact on RBC membrane tension that mediates the reduction in parasite invasion efficiency. We propose that, irrespective of Dantu genotype, there is a tension threshold for successful *P. falciparum* invasion, a novel concept that adds complexity to our understanding of the importance of red cell biomechanical properties^22, 23^ and the heterogeneity of parasite invasion. This model could also explain other well-established features of *P. falciparum* invasion, such as their preference for younger RBCs^24^, which have lower tension and higher radii, and will more often fall beneath the tension threshold for invasion. Several other polymorphisms also affect RBC tension^25, 26^ and further studies will be required to investigate whether the same mechanism might be generalizable across multiple malaria-protective human genetic traits. As well as improving our biological understanding of erythrocyte-parasite dynamics, this study also points the way to the potential for novel malaria interventions based on modifying the biomechanical properties of circulating RBCs.

## Materials and Methods

### Study participants

Samples were obtained from 42 children under the age of 13 years from two cohorts from the Kilifi County on the Indian Ocean coast of Kenya, who were involved in ongoing studies on malaria: (i) 18 children from a cohort subject to annual cross-sectional surveys through which blood samples are collected and frozen, and (ii) 24 children from a cohort recruited at 3-12 months of age and followed up for hospital admission since 2007, whose blood samples were collected and used in the assays within 24 hours of blood draw. Individual written informed consent was provided by the parents of all study participants. Ethical approval for the study was granted by the Kenya Medical Research Institute Scientific and Ethics Review Unit in Nairobi, Kenya (SERU3420 and SERU3500), the NHS Cambridgeshire 4 Research Ethics Committee (REC reference 15/EE/0253), and the Wellcome Sanger Institute Human Materials and Data Management Committee.

### Genotyping Dantu samples

gDNA was extracted from whole blood using a QIAmp 96 DNA QIcube HT kit on a QIAcube HT System (QIAGEN) following manufacturer’s instructions. The restriction fragment length polymorphism (RFLP) assay to detect genotypes at the Dantu marker SNP, rs186873296, has been previously described^4, 5^. Briefly, PCR amplification of the region of interest containing rs186873296 was performed using the following primers: 5’ACGTTGGATGGCAGATTAGCATTCACCCAG3’ and 5’ACGTTGGATGCTCCAGAGTAAGCATCCTTC3’ generating an amplicon of 124bp. Fragmentation of the PCR product was then performed using the CviQI restriction enzyme (NEB), which allowed us to differentiate between non-Dantu homozygotes (AA) that would remain uncut, Dantu heterozygotes (AG) that would generate two bands of 64 and 56bp, and Dantu homozygotes (GG) which would generate a single band of 56bp.

### *In vitro* culture of *P. falciparum* parasites

All *P. falciparum* parasite strains used in this study (3D7, Dd2, SAO75, GB4, 7G8, ΔPfEBA175) were routinely cultured in human O-erythrocytes (NHS Blood and Transplant, Cambridge, UK, and Kenya Medical Research Institute, Nairobi, Kenya) at 3% hematocrit in RPMI 1640 medium with 25 mM Hepes, 20 mM glucose, and 25 μg/mL gentamicin containing 10% Albumax at 37°C (complete medium), under an atmosphere of 1% O_2_, 3% CO_2_, and 96% N_2_ (BOC, Guildford, UK). Parasite cultures were synchronized on early stages with 5% D-sorbitol (Sigma-Aldrich, Dorset, UK). Use of erythrocytes from human donors for *P. falciparum* culture was approved by NHS Cambridgeshire 4 Research Ethics Committee and the Kenya Medical Research Institute Scientific and Ethics Review Unit.

### RBC preference invasion assays

In all cases, blood was collected in EDTA-vacutainers and either used within 24h or cryopreserved using standard methods. Both fresh and frozen/thawed RBCs from Dantu homozygote, heterozygote and non-Dantu children were used in these assays, with no difference in parasite invasion efficiency being observed between them. RBCs were stained with three concentrations of CellTrace Far Red Cell Proliferation kit (Invitrogen, UK) - 1µM, 4µM and 16µM - corresponding to the three genotype groups. After a 2h incubation at 37°C under rotation, the stained RBCs were washed and resuspend to 2% Haematocrit (Hct) with complete medium. The cells were stored at 4°C until use for up to 24h after staining. To evaluate the preference of the parasites to invade RBCs of different Dantu genotype, parasite cultures containing mostly ring forms at 2-5% parasitaemia were pooled with equal volumes of RBCs from each genotype group (25µl pRBCs, 25µl Dantu homozygote RBCs, 25µl Dantu heterozygote RBCs and 25µl non-Dantu RBCs) in the same well in 96-well plates. To evaluate whether the different concentrations of the dye could affect parasite growth, parasite cultures were mixed with stained RBCs from each genotype group in individual wells in a 1:1 ratio (50µl pRBCs + 50µl stained RBCs), while normal parasite growth controls were also evaluated by mixing parasite cultures with unstained RBCs from each genotype group in individual wells in a 1:1 ratio (50µl pRBCs + 50µl unstained RBCs). The samples were incubated for 48h at 37°C under static conditions as described above. After 48h, the cultures were treated with 0.5 mg/mL ribonuclease A (Sigma Aldrich, UK) in PBS for 1h at 37°C to remove any trace of RNA. To evaluate all parasitised RBCs, the cells were stained with 2x SYBR Green I DNA dye (Invitrogen, Paisley, UK) in PBS for 1 h at 37°C. Stained samples were examined with a 488nm blue laser, and a 633nm red laser on a BD FACS Canto flow cytometer (BD Biosciences, Oxford, UK). SYBR Green I was excited by a blue laser and detected by a 530/30 filter. CellTrace Far Red was excited by a red laser and detected by a 660/20 filter. BD FACS Diva software (BD Biosciences, Oxford, UK) was used to collect 50,000 events for each sample. The data collected were then further analysed with FlowJo (Tree Star, Ashland, Oregon) to obtain the percentage of parasitised RBCs within each genotype group. Statistical analyses were performed using R statistical software (version 3.3.3), where differences in invasion across the three Dantu genotype groups were evaluated using a one-way ANOVA test, while pairwise comparisons between genotype groups were evaulatuated using the Tukey HSD test. All experiments were carried out in triplicate and the data are presented as the median and interquartile ranges of invasion data across samples within each genotype group.

### Live invasion imaging

All live imaging assays were performed blind to Dantu genotype group. Highly concentrated (97-100%) infected cells (strain 3D7) were isolated by magnetic separation (LD columns, Miltenyi Biotec, UK) directly before the experiments and re-suspended in complete medium either with Dantu or non-Dantu RBCs at 0.2 % Hct. The Dantu and non-Dantu RBCs suspensions were loaded in separate SecureSeal Hybridization Chambers (Sigma-Aldrich) and imaging was performed at the same time by employing 3 microscopes in order to guarantee the same conditions throughout the experiments. Each sample was recorded for about 2 hours to enable recording of a sufficient number of events. A custom-built temperature control system was used to maintain the optimal culture temperature of 37°C while running these experiments. Samples were placed in contact with a transparent glass heater driven by a PID temperature controller in a feedback loop with the thermocouple attached to the glass slide. A Nikon Eclipse Ti-E inverted microscope (Nikon, Japan) was used with a Nikon 60X Plan Apo VC N.A. 1.40 oil immersion objective, kept at physiological temperature through a heated collar. Motorized functions of the microscope were controlled via custom software written in-house and focus was maintained throughout the experiments using the Nikon Perfect Focus system. Images were acquired in bright-field with red filter using a CMOS camera (model GS3-U3-23S6M-C, Point Grey Research/FLIR Integrated Imaging Solutions (Machine Vision), Ri Inc., Canada) at a frame rate of 4 fps, with pixel size corresponding to 0.0986 µm.

### Optical tweezers

The optical tweezers setup fits onto the same Nikon inverted microscope used for imaging and consists of a solid-state pumped Nd:YAG laser (IRCL-2W-1064; CrystaLaser, Reno, NV) having 2W optical output at a wavelength of 1064 nm. The laser beam was steered via a pair of acousto-optical deflectors (AA Opto-Electronic, Orsay, France) controlled by custom-built electronics that allow multiple trapping with subnanometer position resolution. Videos were taken at 60 frames/s through a 60X Plan Apo VC 1.20 NA water objective (Nikon) with pixel size corresponding to 0.0973 µm. Dantu and non-Dantu RBCs were suspended in complete medium at 0.05 % Hct with purified schizonts and loaded in separate chambers coated with 10 µl solution of poly(l-lysine)-graft-poly(ethylene glycol) (PLL-g-PEG) (SuSoS AG, Dübendorf, Switzerland) at 0.5 mg/mL concentration and incubated for 30 minutes to prevent excessive adherence of cell proteins onto the coverslip. Adhesive forces at the merozoite-erythrocyte interface were quantified by evaluating the elastic morphological response of the erythrocyte as it resisted merozoite detachment^10^. Immediately after schizont egress, merozoites were manipulated by optical trapping and delivered to the surface of uninfected erythrocytes until attachment. Trapping durations were kept short (< 10 seconds) to minimise any possible detrimental effect of local heating because at full laser power, a few degrees Celsius of heating are expected locally around the laser beam focus). Then a second red blood cell was delivered close to the merozoite to form an erythrocyte-merozoite-erythrocyte system^10^. Erythrocyte maximal elongation before detachment was measured by pulling away the erythrocyte that adheres to the merozoite from their point of attachment, while the opposing force, on a second erythrocyte of our system, is given by either a second optical trap or by adhesion to the bottom of the sample chamber. We do not pull on the merozoite directly because this force would be weak and difficult to calibrate. Finally, because erythrocytes are known to behave mechanically as a linear spring in this regime^27^, the merozoite-erythrocyte adhesive forces were calculated by multiplying the erythrocyte end-to-end elongation before detachment and the stiffness of the erythrocyte cell.

### Characterization of RBC membrane by flow cytometry

A panel of antibodies was selected against the 11 antigens that have been confirmed to be or could be potentially involved in cell adhesion and parasite invasion. Each blood sample was diluted at 0.5% haematocrit, washed twice with PBS and incubated in primary mouse monoclonal antibodies for 1h at 37°C. Antibodies used: anti-CD35-APC (CR1, Thermofisher, 1:50); antiCD44-BRIC 222-FITC (1:100, IBGRL); Integrin: anti-CD49d-APC (1:50, Milteny Biotec); anti-CD55-BRIC-216-FITC (1:500, IBGRL); Transferrin R: anti-CD71-FITC (1:100, ThermoFisher); Basigin: anti-CD147-FITC (1:100, ThermoFisher); Band3: anti-CD233-BRIC6-FITC (1:1000, IBGRL); Duffy antigen: anti-CD234-APC (1:100, Milteny Biotec); GYPA: CD235a-BRIC 256-FITC (1:1000, IBGRL); GYPC: anti-CD236R-BRIC10-FITC (1:1000, IBGRL).

For detection of GYPB, first cells were incubated with an anti-GYPB (1:100, rabbit polyclonal antibody, Abcam), then washed twice with PBS and then incubated with a goat-anti-rabbit AlexFluor488 labelled antibody. After incubation, cells were washed twice in PBS and analyzed on a BD FACS Canto flow cytometer. Data were analyzed using FlowJo Software (Treestar, Ashland, Oregon). Statistical analyses to test differences in RBC membrane surface expression across genotype groups were performed using R statistical software (version 3.3.3).

### Erythrocyte plasma membrane profiling

Plasma membrane profiling was performed as previously described^14, 15^. Briefly, three of each Dantu genotype RBC samples were washed with PBS. Surface sialic acid residues were oxidized with sodium meta-periodate (Thermo) then biotinylated with aminooxy-biotin (Biotium). After quenching, cells were incubated in 1% Triton X-100 lysis buffer. Biotinylated glycoproteins were enriched with high affinity streptavidin agarose beads (Pierce) and washed extensively. Captured protein was denatured with DTT, alkylated with iodoacetamide (IAA, Sigma) and digested on-bead with trypsin (Promega) in 200 mM HEPES pH 8.5 for 3h. Tryptic peptides were collected and labelled using TMT reagents. The reaction was quenched with hydroxylamine, and TMT-labelled samples combined in a 1:1:1:1:1:1:1:1:1 ratio. Labelled peptides were enriched, desalted, and 80% of the combined sample separated into twelve fractions using high pH reversed phase HPLC as previously described^28^. 100% of six fractions in addition to 50% of the remaining unfractionated sample were subjected to mass spectrometry.

Mass spectrometry data were acquired using an Orbitrap Fusion Lumos (Thermo Fisher Scientific, San Jose, CA) interfaced via an EASYspray source to an Ultimate 3000 RSLC nano UHPLC. Peptides were loaded onto a 100 µm ID x 2 cm Acclaim PepMap nanoViper precolumn (Thermo Fisher Scientific) and resolved using a 75 µm ID x 50 cm 2 µm particle PepMap RSLC C18 EASYspray column. Loading solvent was 0.1% FA, analytical solvent A: 0.1% FA and B: 80% MeCN + 0.1% FA. All separations were carried out at 40°C. Samples were loaded at 5 µL/minute for 5 minutes in loading solvent before beginning the analytical gradient. The following gradient was used: 3-7% solvent B over 2 minutes, 7-37% solvent B over 173 minutes, followed by a 4 minute wash at 95% solvent B and equilibration at 3% solvent B for 15 minutes. Each analysis used a MultiNotch MS3-based TMT method^29^. The following settings were used: MS1: 380-1500 Th, 120,000 Resolution, 2×10^5^ automatic gain control (AGC) target, 50 ms maximum injection time. MS2: Quadrupole isolation at an isolation width of m/z 0.7, CID fragmentation (normalised collision energy (NCE) 35) with ion trap scanning in turbo mode from m/z 120, 1.5×10^4^ AGC target, 120 ms maximum injection time. MS3: In Synchronous Precursor Selection mode the top 10 MS2 ions were selected for HCD fragmentation (NCE 65) and scanned in the Orbitrap at 60,000 resolution with an AGC target of 1×10^5^ and a maximum accumulation time of 150 ms. Ions were not accumulated for all parallelisable time. The entire MS/MS/MS cycle had a target time of 3 s. Dynamic exclusion was set to +/- 10 ppm for 70 s. MS2 fragmentation was trigged on precursors 5×10^3^ counts and above.

### Mass spectrometry data analysis

Mass spectra were processed using a Sequest-based software pipeline for quantitative proteomics, “MassPike”, through a collaborative arrangement with Professor Steven Gygi’s laboratory at Harvard Medical School. MS spectra were converted to mzXML using an extractor built upon Thermo Fisher’s RAW File Reader library (version 4.0.26). In this extractor, the standard mzxml format has been augmented with additional custom fields that are specific to ion trap and Orbitrap mass spectrometry and essential for TMT quantitation. These additional fields include ion injection times for each scan, Fourier Transform-derived baseline and noise values calculated for every Orbitrap scan, isolation widths for each scan type, scan event numbers, and elapsed scan times. This software is a component of the MassPike software platform and is licensed by Harvard Medical School.

A combined database was constructed from the human Uniprot database (26th January, 2017) and common contaminants such as porcine trypsin and endoproteinase LysC. The combined database was concatenated with a reverse database composed of all protein sequences in reversed order. Searches were performed using a 20 ppm precursor ion tolerance. Fragment ion tolerance was set to 1.0 Th. TMT tags on lysine residues and peptide N termini (229.162932 Da) and carbamidomethylation of cysteine residues (57.02146 Da) were set as static modifications, while oxidation of methionine residues (15.99492 Da) was set as a variable modification.

To control the fraction of erroneous protein identifications, a target-decoy strategy was employed^30^. Peptide spectral matches (PSMs) were filtered to an initial peptide-level false discovery rate (FDR) of 1% with subsequent filtering to attain a final protein-level FDR of 1%. PSM filtering was performed using a linear discriminant analysis, as described previously^30^. This distinguishes correct from incorrect peptide IDs in a manner analogous to the widely used Percolator algorithm (https://noble.gs.washington.edu/proj/percolator/), though employing a distinct machine learning algorithm. The following parameters were considered: XCorr, ΔCn, missed cleavages, peptide length, charge state, and precursor mass accuracy.

Protein assembly was guided by principles of parsimony to produce the smallest set of proteins necessary to account for all observed peptides (algorithm described in^30^. Proteins were quantified by summing TMT reporter ion counts across all matching peptide-spectral matches using “MassPike”, as described previously^29^. Briefly, a 0.003 Th window around the theoretical m/z of each reporter ion (126, 127n, 127c, 128n, 128c, 129n, 129c, 130n, 130c, 131n, 131c) was scanned for ions, and the maximum intensity nearest to the theoretical m/z was used. The primary determinant of quantitation quality is the number of TMT reporter ions detected in each MS3 spectrum, which is directly proportional to the signal-to-noise (S:N) ratio observed for each ion. Conservatively, every individual peptide used for quantitation was required to contribute sufficient TMT reporter ions (minimum of ~1250 per spectrum) so that each on its own could be expected to provide a representative picture of relative protein abundance^29^. An isolation specificity filter with a cutoff of 50% was additionally employed to minimise peptide co-isolation^29^. Peptide-spectral matches with poor quality MS3 spectra (more than 8 TMT channels missing and/or a combined S:N ratio of less than 250 across all TMT reporter ions) or no MS3 spectra at all were excluded from quantitation. Peptides meeting the stated criteria for reliable quantitation were then summed by parent protein, in effect weighting the contributions of individual peptides to the total protein signal based on their individual TMT reporter ion yields. Protein quantitation values were exported for further analysis in Excel.

Proteins were filtered to include those most likely to be present at the cell surface with high confidence. These comprised proteins with Uniprot Subcellular Location (www.uniprot.org) terms matching ‘Multipass’, ‘GPI anchored’, ‘Lipid Anchored’, ‘Type I transmembrane’, ‘Type II transmembrane’, ‘Type III transmembrane’, ‘Type IV transmembrane’, and those predicted to have transmembrane regions based on TMHMM version 2.0^31^.

For protein quantitation, reverse and contaminant proteins were removed. Despite extensive washing of biotinylated proteins when bound to Streptavidin beads, variable levels of contamination with abundant haemoglobin components were nevertheless detectable. As opposed to normalisation assuming equal protein loading across all channels, normalisation was instead performed from the summed signal:noise values of all proteins passing the filter described above. For further analysis and display in figures, only these filtered proteins are displayed. For **Fig. 2c** and **Supplementary Fig. 3b**, fractional TMT signals were used (i.e. reporting the fraction of maximal signal observed for each protein in each TMT channel). For **Fig. 2b**, fold change was calculated on the basis of (average signal:noise (Dantu homozygotes) / average signal:noise (non-Dantu). For figure **Supplementary Fig. 3a**, fold change was calculated for each Dantu variant donor by (signal:noise (Dantu variant) / average signal:noise (non-Dantu).

For **Fig. 2a**, the method of significance A was used to estimate the p-value that each protein ratio was significantly different to 1. Values were calculated and corrected for multiple hypothesis testing using the method of Benjamini-Hochberg in Perseus version 1.5.1.6^32^. For **Supplementary Fig. 3b**, two-tailed Student’s t-test values were calculated and corrected for multiple hypothesis testing using the method of Benjamini-Hochberg in Excel. Hierarchical centroid clustering based on uncentered correlation was performed using Cluster 3.0 (Stanford University) and visualised using Java Treeview (http://jtreeview.sourceforge.net).

### Erythrocyte membrane contour detection and flickering spectrometry

Dantu and non-Dantu RBCs were diluted into culture medium at 0.01 % Hct and loaded in different chambers to provide an optimal cell density and avoid overlapping cells. All live-cell experiments were performed at 37°C by using the setup described in the previous section. 20 seconds time-lapse videos were recorded at high frame rate 514 frames/s and short exposure time 0.8 ms. The contour of the erythrocyte membrane was detected with subpixel resolution by an optimised algorithm developed in house and implemented in Matlab (The MathWorks, Natick, MA). Detailed explanation of the membrane fluctuation analysis has been described in previous work^18^.

## Data Availability

The authors declare that the data supporting the findings of this study are available within the manuscript and its Supplementary information files.

## Supporting information

## Acknowledgements

We thank Emmanuel Mabibo, Jacob Golijo, Alphonse Kazungu, Ruth Mwarabu, the staff of Kilifi County Hospital and the KEMRI-Wellcome Trust Research Programme, Kilifi, for their help with participant recruitment, data and sample collection, and to Ellen Leffler and Gavin Band for discussions on the study. We also thank the study participants and their parents for agreeing to participate in this study. JCR, AM and DK were supported by the Wellcome Trust (098051). MPW is funded by a Wellcome Senior Fellowship (108070). TNW is funded through Fellowships awarded by the Wellcome Trust (091758 and 202800). SNK is supported by the Wellcome Trust-funded Initiative to Develop African Research Leaders (IDeAL) early-career postdoctoral fellowship. The Wellcome Trust provides core support to The KEMRI/Wellcome Trust Research Programme, Kilifi, Kenya (084535), Wellcome Sanger Institute, Cambridge, UK (206194/Z/17/Z) and the Wellcome Centre for Human Genetics, Oxford, UK (090532/Z/09/Z, 203141). PC is supported by the Engineering and Physical Sciences Research Council (EPSRC) (<u>EP/R011443/1</u>), and VI is supported by the EPSRC and the Sackler fellowship. This paper is published with permission from the Director of KEMRI.

