## Supplementary material for "Red blood cell tension controls *Plasmodium falciparum* invasion and protects against severe malaria in the Dantu blood group"

**Supplementary Table 1 | Study participant genotypes at 6 malaria-protective polymorphisms**

| Sample | Dantu (rs186873296) | *HbS* (rs334) | Alpha thalassaemia | *G6PD* (rs1050828) | *ATP2B4* (rs1541255) | *CR1* (rs17047661) | *ABO* (rs8176719) |
|---|---|---|---|---|---|---|---|
| MSC1 | GG | AA | Het | CC | AA | AG | DI |
| MSC2 | AG | AA | Norm | CT | AA | AA | DD |
| MSC3 | AA | AA | Norm | CC | AA | AG | DD |
| MSC4 | GG | AA | Norm | TT | AG | AG | II |
| MSC5 | AG | AA | Norm | CC | AA | AG | DD |
| MSC6 | AA | AA | Norm | CC | AA | - | - |
| MSC7 | GG | AA | Norm | - | - | - | - |
| MSC8 | AG | AA | Norm | CC | AA | AG | DD |
| MSC9 | AA | AA | Norm | CC | AA | GG | DI |
| MSC 10 | GG | AA | Het | - | - | - | - |
| MSC 11 | AG | AA | Het | CC | AA | AG | DD |
| MSC 12 | AA | AA | Norm | TT | AA | - | - |
| MSC 13 | GG | AA | Het | - | - | - | - |
| MSC 14 | AG | AA | Het | TT | AA | GG | DI |
| MSC 15 | AA | AA | Norm | CC | AA | GG | DI |
| MSC 16 | GG | - | - | - | - | - | - |
| MSC 17 | AG | AA | Het | CT | AA | AG | DI |
| MSC 18 | AA | AA | Norm | CC | AA | AG | DD |
| KBC1 | GG | AA | Norm | CC | AA | GG | DI |
| KBC2 | AA | AA | Norm | CC | AA | GG | DI |
| KBC3 | AG | AA | Norm | CC | AA | GG | DI |
| KBC4 | GG | AA | Norm | CC | GG | AG | DD |
| KBC5 | AA | AA | Norm | CC | GG | AG | DD |
| KBC6 | AA | AA | Norm | CC | AA | GG | DD |
| KBC7 | GG | AA | Norm | CC | AA | GG | DD |
| KBC 8 | AG | AA | Norm | CC | AA | GG | DD |
| KBC 9 | AG | AA | Norm | CC | AG | GG | DI |
| KBC 10 | GG | AA | Norm | CC | AG | GG | DI |
| KBC 11 | AA | AA | Norm | CC | AG | GG | DI |
| KBC 12 | AG | AA | Norm | CC | AA | AG | DD |
| KBC 13 | GG | AA | Norm | CC | AA | AG | DD |
| KBC 14 | AA | AA | Norm | CC | AA | AG | DD |
| KBC 15 | AG | AA | Norm | CC | AA | GG | DI |
| KBC 16 | AA | AA | Norm | CC | AA | GG | II |
| KBC 17 | GG | AA | Norm | CC | AA | GG | II |
| KBC 18 | AA | AA | Norm | CC | AG | GG | DD |

| | | | | | | | |
|---|---|---|---|---|---|---|---|
| KBC 19 | AG | AA | Norm | CC | AG | GG | DD |
| KBC 20 | GG | AA | Norm | CC | AG | GG | DD |
| KBC 21 | AA | AA | Norm | CC | AA | GG | DI |
| KBC 22 | AG | AA | Norm | CC | GG | AG | DI |
| KBC 23 | AA | AA | Norm | CC | GG | AG | DI |
| KBC 24 | AG | AA | Norm | CC | AA | GG | II |

Study participant genotype information for malaria-protective polymorphisms including Dantu (rs186873296), *HBB* (rs334), alpha-thalassaemia, *G6PD* (rs1050828), *ATP2B4* (rs1541255) and *CR1* (rs17047661), and *ABO* (rs8176719) – a key SNP in determining blood group O status, where two copies of a deletion at this site (DD) indicates blood group type O, while one copy of the deletion (DI) or no deletion (II) at this site indicates blood type A, B or AB. Study subjects were obtained from two cohorts, from which both frozen and fresh red blood cell (RBC) samples were obtained. Genotypes were controlled within each experimental set of Dantu homozygote (rs186873296=GG), heterozygote (rs186873296=AG), and non-Dantu (rs186873296=AA) individuals.

**Supplementary Table 2 | Preference invasion into Dantu variant RBCs**

| | **one-way ANOVA across Dantu genotype groups** | **Pairwise comparisons of mean parasitaemia differences across Dantu genotype groups** | | | | | |
|---|---|---|---|---|---|---|---|
| **Parasite strain** | **P value** | **non-Dantu-Dantu hom (%)** | **P adj** | **non-Dantu-Dantu het (%)** | **P adj** | **Dantu het-Dantu hom (%)** | **P adj** |
| 3D7 | 0.001 | 2.1134 (0.9551 - 3.2718) | 0.001 ** | 0.9569 (-0.2014 - 2.1153) | 0.114 | 1.1565 (-0.0019 - 2.3149) | 0.050 |
| Dd2 | 0.018 | 2.7888 (0.535 - 5.0426) | 0.015 * | 1.012 (-1.2418 - 3.2658) | 0.490 | 1.7768 (-0.477 - 4.0306) | 0.135 |
| SAO75 | 0.035 | 2.0798 (0.212 - 3.9476) | 0.028 * | 0.8854 (-0.9824 - 2.7532) | 0.454 | 1.1945 (-0.6733 - 3.0623) | 0.252 |
| GB4 | 0.122 | 0.6852 (-1.6704 - 3.0408) | 0.719 | 1.8461 (-0.3747 - 4.067) | 0.107 | 1.1609 (-1.1947 - 3.5166) | 0.961 |
| 7G8 | 0.212 | 0.9928 (-1.7233 - 3.7089) | 0.619 | 1.9408 (-0.7753 - 4.6569) | 0.186 | 0.948 (-1.7681 - 3.6641) | 0.967 |
| ∆PfEBA175 | 0.028 | 1.3932 (0.056 - 2.7303) | 0.040 * | 1.2441 (-0.093 - 2.5813) | 0.072 | 0.1491 (-1.2146 - 1.5127) | 0.408 |
| ∆PfEBA140 | 0.448 | 0.8471 (-1.3268 - 3.0209) | 0.610 | 1.072 (-1.1019 - 3.2459) | 0.457 | 0.2249 (-1.992 - 2.4418) | 0.645 |

Mean difference in parasitaemia between Dantu genotype groups (%) is compared using the one-way ANOVA, and pairwise comparisons using Tukey HSD test. The lower and upper limits of the differences in parasitaemia are given in parentheses. P adj – p-value after adjustment for the multiple pairwise comparisons with Benjamini-Hochberg FDR. ** $p < 0.01$; * $p < 0.05$

**Supplementary Table 3 | Biomechanical properties of RBCs**

**(a)**

| | Non-Dantu (Mean ± SD) | Dantu heterozygote (Mean ± SD) | Dantu homozygote (Mean ± SD) |
|---|---|---|---|
| Number of cells | 249 | 252 | 247 |
| Bending modulus ($10^{-20}$ J) | 14.0 ± 1.5 | 14.0 ± 1.8 | 13.0 ± 2.7 |
| Tension ($10^{-7}$ N/m) | 6.0 ± 1.9 | 7.9 ± 2.8 | 8.8 ± 0.7 |
| Radius (µm) | 4.2 ± 0.1 | 4.0 ± 0.1 | 3.9 ± 0.1 |
| Viscosity ($10^{-3}$ Pa s) | 20.5 ± 5.6 | 20.8 ± 5.3 | 23.5± 4.6 |

**(b)**

| | Non-Dantu (Mean ± SD) | Dantu heterozygote (Mean ± SD) | Dantu homozygote (Mean ± SD) |
|---|---|---|---|
| Number of donors | 6 | 6 | 6 |
| Bending modulus ($10^{-20}$ J) | 14.0 ± 1.5 | 14.0 ± 1.8 | 13.0 ± 2.7 |
| Tension ($10^{-7}$ N/m) | 5.7 ± 1.9 | 7.9 ± 2.8 | 8.7 ± 0.7 |
| Radius (µm) | 4.2 ± 0.1 | 4.0 ± 0.1 | 3.9 ± 0.1 |
| Viscosity ($10^{-3}$ Pa s) | 20.5 ± 5.6 | 20.8 ± 5.3 | 23.5± 4.6 |

The membrane mechanics of RBCs, measured with live video flickering spectrometry, were compared across Dantu genotypes. **(a)** Mean and standard deviation of the distribution of all cells. **(b)** Mean and standard deviation of averages from each donor. Fits for bending and tension are for $7^{th}$ to $20^{th}$ q (wave vector) modes; fits for viscosity for $7^{th}$ to $11^{th}$ q modes. Data shown in Figure 3a.

**Supplementary Table 4 | The impact of RBC tension on membrane deformation and parasite invasion**

| **Invasion efficiency** | **Genotype group** | **Number of cells** | **Tension (10$^{-7}$ N/m)** (Mean ± SD) | **Deformation score** |
|---|---|---|---|---|
| Successful | non-Dantu | 44 | 1.7 ± 1.2 | 0/1 |
| | Dantu homozygotes | 31 | 2.6 ± 2.2 | 0/1 |
| Failed | non-Dantu | 40 | 8.8 ± 5.7 | 2/3 |
| | Dantu homozygotes | 48 | 9.4 ± 4.9 | 2/3 |

The impact of tension on membrane deformation induced by parasites during pre-invasion phase, and on subsequent invasion, was compared across Dantu genotypes. Mean and Standard deviation is reported for tension. Data shown in Figure 3c, d.

**Supplementary Table 5 | Merozoite-RBC adhesion force measured by optical tweezers**

| | Non-Dantu (Mean ± SD) | Dantu heterozygote (Mean ± SD) | Dantu homozygote (Mean ± SD) |
|---|---|---|---|
| Number of cells | 19 | 21 | 24 |
| Merozoite-erythrocyte attachment force | 42.5 ± 15.7 | 49.5 ± 22.9 | 39.8 ± 15.6 |

The strength of attachment between merozoites and Dantu RBCs was measured using optical tweezers, where adhesive forces at the merozoite-RBC interface were quantified and compared across genotype groups, by evaluating the elastic morphological response of the erythrocyte as it resisted merozoite detachment. Data shown in Supplementary Figure 2.

**Supplementary Figure 1 | RBC membrane deformation scores triggered by merozoite contact**

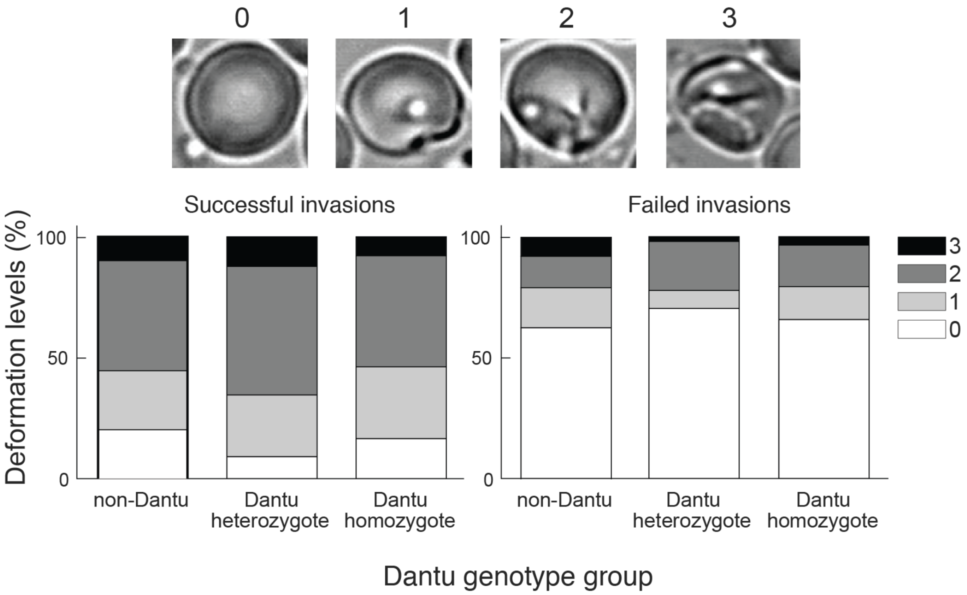

The degree to which merozoites deformed individual erythrocytes was given by a simplified four-point deformation scale (0, 1, 2, and 3), based on the most extreme degree of deformation achieved (Weiss, G. E. *et al*. *PLoS Pathog* 2015). The degree of deformation was compared across parasites that successfully invaded RBCs ("Invaders") and parasites that did not invade ("Non-invaders"). Deformation was also compared across Dantu genotype groups.

**Supplementary Figure 2 | Merozoite-RBC adhesion force**

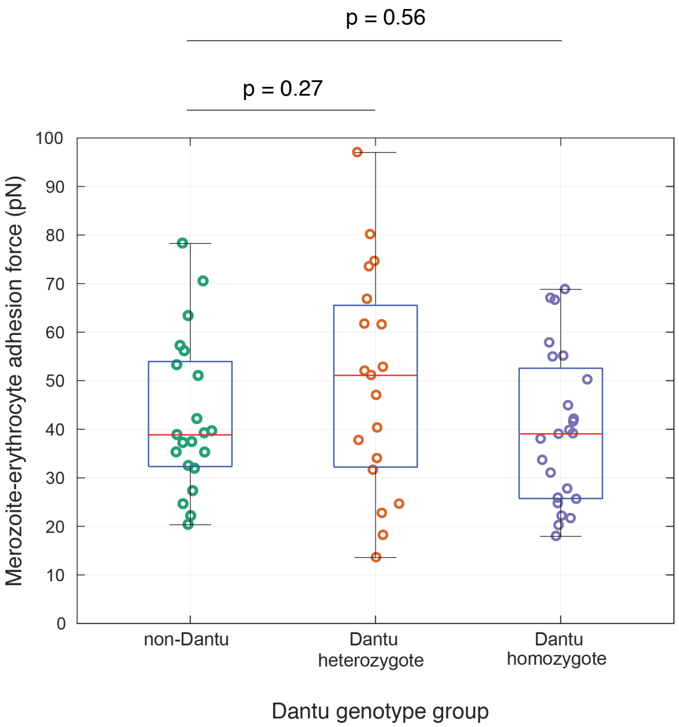

Merozoite adhesion force to Dantu RBCs was measured and compared across genotype groups using optical tweezers. Merozoites attached to RBCs were pulled using optical traps, and the adhesive forces at the merozoite-RBC interface were quantified by evaluating the elastic morphological response of the RBC as it resisted merozoite detachment. Data in Supplementary Table 5. Six RBC samples per genotype group were tested. The median is indicated by the middle red line in the boxplots, with the 25$^{th}$ and 75$^{th}$ percentiles indicated by the tops and bottoms of each plot, while whiskers denote total data range. If the median is not centered in the box, it shows sample skewness.

**Supplementary Figure 3 |** Plasma membrane profiling by tandem mass tag (TMT)-based MS3 mass spectrometry

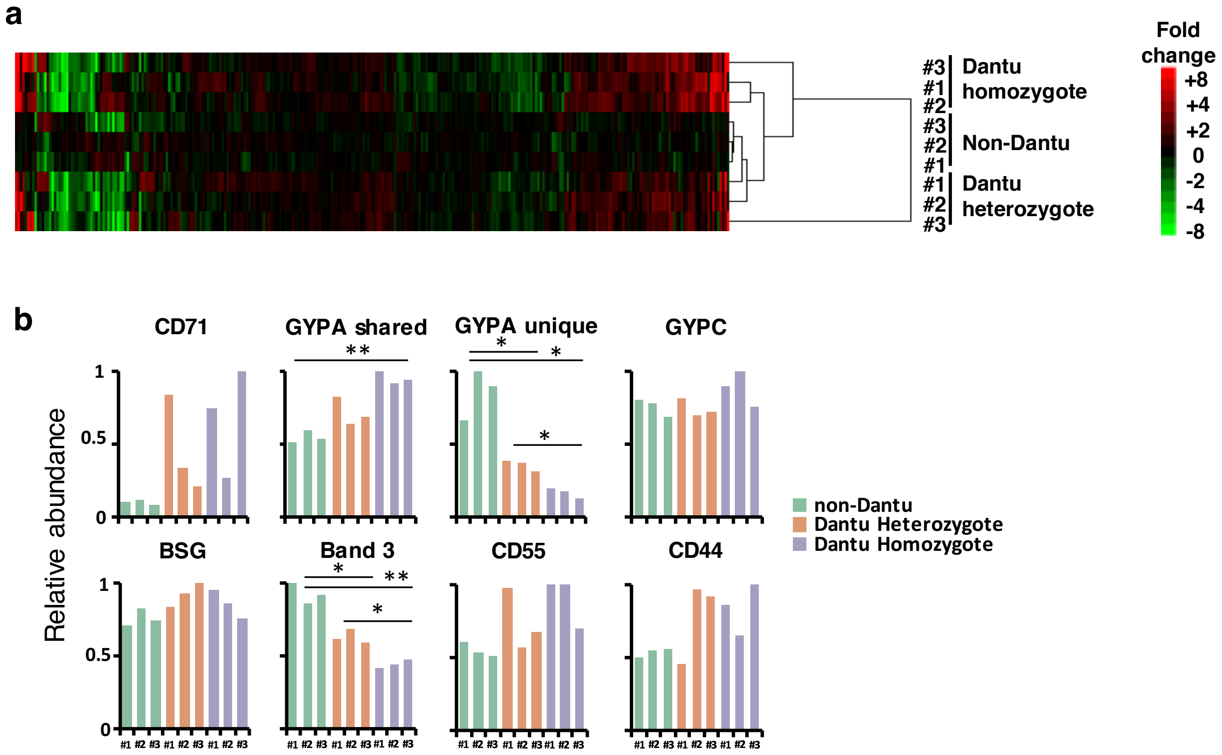

The impact of the Dantu polymorphism on RBC membrane protein expression levels was quantified using mass spectrometry. (**a**) Hierarchical cluster analysis of all proteins quantified and annotated as described in the Methods. Fold change was calculated for each donor by (signal:noise (donor) / average signal:noise (non-Dantu). (**b**) Proteomic quantification of markers shown in **Figure 2a**. All markers were quantified by proteomics apart from GYPB. Statistical comparisons of quantitative protein expression across Dantu genotype groups were performed using two-tailed t-test with multiple hypothesis correction: * $p < 0.05$, ** $p < 0.01$.

**Supplementary Figure 4 |** Parasite preference invasion measured by flow cytometry

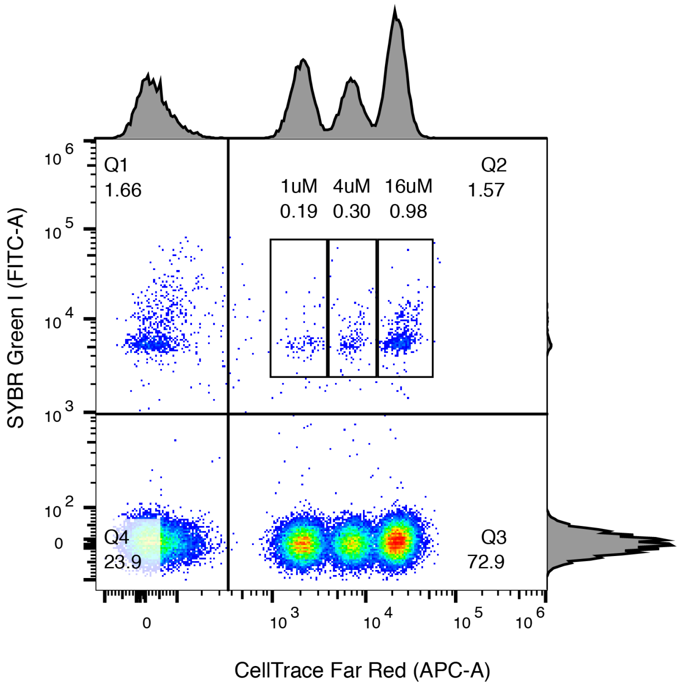

Parasite invasion preference for RBCs from donors across Dantu genotype groups (Dantu homozygous, Dantu heterozygous and non-Dantu) was measured using a flow cytometry-based preference invasion assay. RBCs from the three Dantu genotypes were differentially labelled with three concentrations of a fluorescent cytoplasmic dye, CellTrace Far Red (x-axis), distinguishing the three RBC populations, while parasite-infected RBCs were detected with a fluorescent DNA dye, SYBR Green I (y-axis). The dot plot represents preference invasion data from 3D7 parasite strain generated by flow cytometry, with the Dantu homozygous, heterozygous and non-Dantu RBCs labelled with 1uM, 4uM and 16uM CellTrace Far Red, respectively. The four distinguishable populations in the dot plot are: unlabeled, infected RBCs (upper left panel, “Q1”); labeled, infected RBCs (upper right panel, “Q2”); labeled, uninfected RBCs (lower right panel, “Q3”); and unlabeled, uninfected RBCs (lower left panel, “Q4”). Gates are drawn around the three clusters of labeled parasitised RBCs in Q2, with the percentage of parasitised RBCs in each cluster indicated.
