## Supplementary material for "Red blood cell tension controls *Plasmodium falciparum* invasion and protects against severe malaria in the Dantu blood group"

**Supplementary Table 6 | Proteomic data from plasma membrane profiling by tandem mass tag (TMT)-based MS3 mass spectrometry**
(A) Spreadsheet of all proteomic data from Figures 2b, 2c and Supplementary Figure 3. All 295 proteins either anchored in the RBC membrane or having a transmembrane region. Signal:noise values were normalised as described in Methods. (B) Data for all quantified proteins without normalisation.

### Videos:

**Supplementary Video 1 | Merozoite invasion studied by time-lapse video recordings in non-Dantu RBCs.** Merozoite invasion into non-Dantu RBCs was studied using time-lapse live video microscopy (4 frames/s), enabling the evaluation of the merozoite invasion efficiency as well as the kinetics of the entire invasion process. Merozoites performed a complex multistep process to invade RBCs in less than one minute: merozoite-erythrocyte initial reversible contacts triggered RBC membrane deformations, and were rapidly followed by merozoite reorientation. After achieving an apical alignment perpendicular to the RBC membrane, the merozoite penetrated the RBC until its complete internalization. Invasion was followed by a reversible morphological change called echinocytosis and finally a ring was formed.

**Supplementary Video 2 | Merozoite invasion studied by time-lapse video recordings in Dantu homozygote RBCs.** Merozoite invasion into Dantu variant RBCs was studied using time-lapse live video microscopy (4 frames/s). Merozoites contacted and deformed Dantu RBC membranes many times in different points of the RBC surface without proceeding to invasion.

**Supplementary Video 3 | Merozoite-erythrocyte adhesion force measured by optical tweezers.**
An erythrocyte-merozoite-erythrocyte bridge was formed by trapping and moving an erythrocyte onto a nearby erythrocyte undergoing invasion. One optical trap was used to keep one erythrocyte fixed, while another trap pulled the second erythrocyte in a normal direction away from the point of merozoite attachment, until detachment.
Adhesive forces at the merozoite-erythrocyte contact were quantified by measuring the maximum elongation of erythrocyte before detachment, as described in Materials and Methods section.

**Supplementary Video 4 | Video recording of all RBC membrane fluctuations around a schizont before egress used to determine their membrane tension.**
Videos at 514 frames/s were recorded of healthy RBCs around a schizont a few minutes before its egress. At this high frame rate, it is possible to detect each RBC contour and analyse their membrane fluctuations to obtain

biophysical parameters such as tension, bending modulus, radius, and viscosity without altering or interfering with RBCs.

**Supplementary Video 5| Egress-invasion process following Supplementary Video 4.**
After measuring tension and the other biophysical properties for all RBCs near the schizont, we recorded schizont egress and the successive invasion process by the newly released merozoites. We then correlated the RBC biomechanical characteristics with their aptitude to be successfully invaded or not.
